# A bispecific antibody targeting a membrane-proximal ASGPR1 region promotes TNF-α recruitment and uptake

**DOI:** 10.64898/2026.09.11.750848

**Authors:** Xu Song, SiJie Li, Aojiasi Hashan, JianMing Du, YiTong Sun, XiaoTao Jiang, Xuan Liu

## Abstract

Recruiting an endocytic receptor through an antibody offers a route to redirect extracellular cytokines into cells. Here, we developed BiAb-J31, an IgG-like bispecific antibody linking TNF-α recognition to a membrane-proximal region of asialoglycoprotein receptor 1 (ASGPR1). J31 was isolated by immunization with an engineered ASGPR1 extracellular domain followed by screening against wild-type ASGPR1 and HepG2 cells. Its variable domains were combined with unmodified adalimumab variable regions using knobs-into-holes and CrossMab engineering, with L234A/L235A/P329G Fc substitutions. BiAb-J31 retained HepG2 binding and engaged both antigens in bridging assays. Sequential binding experiments demonstrated recruitment of TNF-α to BiAb-J31-treated cells. Live-cell imaging showed uptake of fluorescent antibody-containing complexes and overlap with LysoTracker-positive compartments. In HepG2 cultures containing BiAb-J31, TNF-α and a PE-conjugated detection antibody, supernatant PE fluorescence fell by approximately 60% over 90 minutes. Truncation and synthetic-peptide assays localized J31 recognition to ASGPR1 residues 62–100, outside the carbohydrate-recognition domain, and AlphaFold 3 modeling proposed an Fv–peptide interface. These findings identify a membrane-proximal ASGPR1-binding antibody that is compatible with bispecific engineering and supports cellular recruitment and uptake of TNF-α-containing complexes. BiAb-J31 provides a molecular starting point for antibody-based cytokine-redirecting approaches.

## 1 Introduction

TNF-α is a major therapeutic target in inflammatory disease. Anti-TNF antibodies established that selective cytokine inhibition can alter disease activity, and adalimumab has demonstrated clinical benefit in rheumatoid arthritis ^[1–3]^. TNF antagonists are also central to inflammatory bowel disease treatment, although primary non-response, loss of response and differences associated with treatment sequencing remain clinically relevant ^[4–6]^. These observations motivate investigation of additional ways to control extracellular TNF-α. One approach is to couple cytokine recognition to cellular uptake, adding a route for cellular internalization alongside the binding activity of an antibody. Such an approach could provide sustained cytokine depletion beyond the reversible binding achieved by neutralizing antibodies and may address TNF-α pools that are less accessible to conventional antagonism.

Targeted protein degradation has expanded the range of strategies for controlling protein abundance ^[7,8]^. Lysosome-targeting chimaeras (LYTACs) extend this concept to extracellular and membrane-associated proteins by linking a target-binding component to an internalizing receptor. The initial CI-M6PR-directed LYTACs established this principle ^[9]^, followed by GalNAc-based constructs recruiting the asialoglycoprotein receptor (ASGPR) ^[10,11]^. ASGPR-directed MoDE-As and heterobifunctional PCSK9 ligands further demonstrated removal of soluble proteins, with the latter study including bispecific antibody formats ^[12,13]^. This expanding range of extracellular degraders provides a framework for receptor-directed cytokine handling ^[14]^.

ASGPR is attractive for this purpose because it participates in hepatic recognition and endocytosis of circulating glycoproteins ^[15–17]^. Its physiological roles include coupling recognition of desialylated platelets to thrombopoietin production ^[18]^. The structurally characterized carbohydrate-recognition domain provides the binding site exploited by glycan-based recruitment strategies ^[19]^. Antibodies recognizing other receptor regions offer alternative attachment sites and molecular geometries. Designed EndoTags have shown that protein binders can recruit ASGPR and other trafficking receptors through sites distinct from those engaged by native ligands ^[20]^; however, EndoTags are de novo designed proteins rather than antibodies, and their cargo-binding specificity requires separate engineering. The specific receptor-binding region and its compatibility with a cargo-binding format are therefore important features of a new recruiter.

Bispecific antibodies provide a genetically encoded framework for connecting these activities ^[21]^. Control of heavy- and light-chain association, developed through heterodimeric IgG engineering, knobs-into-holes and immunoglobulin domain crossover, makes it possible to combine two binding specificities in an IgG-like molecule ^[22–25]^. For cytokine uptake, this architecture allows a defined cytokine-binding arm to be paired with an independently selected receptor-binding antibody.

Here, we describe J31, an antibody recognizing the membrane-proximal 62–100 region of ASGPR1, and its incorporation into the ASGPR1×TNF-α bispecific antibody BiAb-J31. We obtained J31 using an immunogen in which residues 237–266 — a segment overlapping the carbohydrate-recognition domain — were replaced with a flexible linker, redirecting the immune response toward other extracellular regions. Selection then used wild-type receptor protein and HepG2 cells. We combined J31 with unmodified adalimumab variable regions and evaluated dual-antigen engagement, cell-associated TNF-α recruitment and uptake in fluorescent detection-antibody systems. Regional mapping and structural modeling define the molecular feature underlying this alternative ASGPR1 attachment site. The study connects a mapped receptor-binding antibody to a cytokine-recruitment application and establishes a basis for further development of extracellular TNF-α redirection.

## 2 Results

### 2.1 An engineered ASGPR1 immunogen yields antibodies recognizing the wild-type receptor

To obtain antibodies recognizing ASGPR1 outside the carbohydrate-recognition domain, we replaced residues Arg237–Asp266 within the extracellular domain with a (G4S)3 linker. This segment overlaps the structurally characterized CRD ^[19]^, and its replacement was intended to reduce the dominance of CRD-directed responses in the immunogen. The resulting immunogen comprised residues 62–291 with the indicated replacement and a C-terminal 10×His tag (Fig. 1A,B). Expression supernatant and nickel-affinity-purified material were examined by SDS-PAGE (Fig. 1C,D).

**Figure 1.**
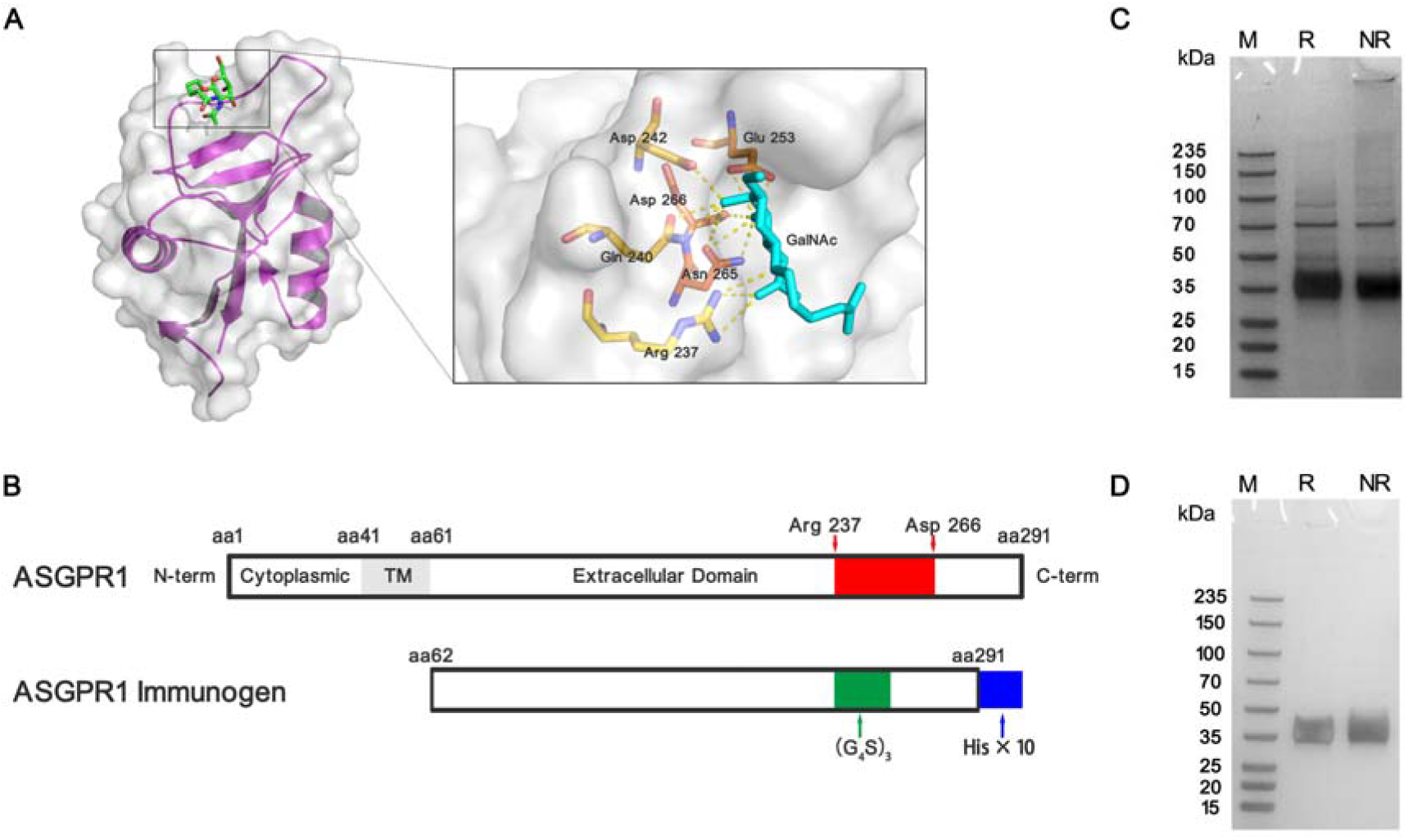
Engineering and preparation of the ASGPR1 immunogen. (A) Structural context of the ASGPR1 carbohydrate-binding region. PDB entry 6YAU contains the GalNAc-derived ligand GN-A. (B) Immunogen design: ASGPR1 residues 62–291 with Arg237–Asp266 replaced by (G4S)3 and a C-terminal 10×His tag. (C) SDS-PAGE of expression supernatant. (D) SDS-PAGE of the nickel-affinity-purified sample.

Mice were immunized on days 0, 14 and 28, received an intravenous boost on day 42, and provided splenocytes for NS-1 fusion on day 45. Screening first used wild-type ASGPR1 extracellular protein in ELISA and then HepG2 immunofluorescence, linking recognition of the unmodified antigen to cell binding (Fig. 2). Seven hybridoma clones, J31–J37, were recovered. All seven bound HepG2 cells in the displayed images. All seven clones also bound Huh7 cells, while HEK293T cells showed no detectable signal, consistent with ASGPR1-dependent recognition.

**Figure 2.**
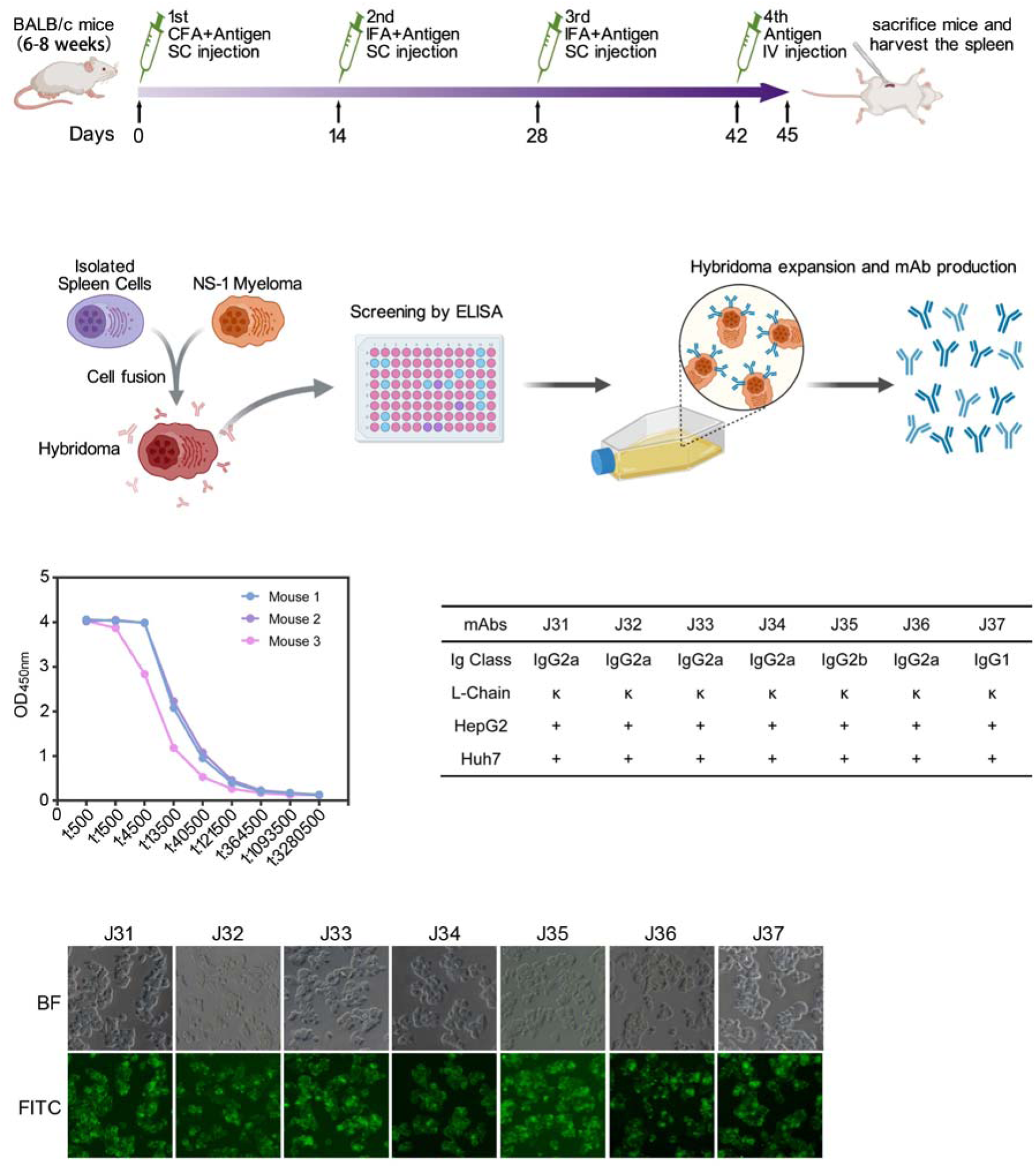
Generation and screening of ASGPR1-binding hybridomas. The upper schematic shows subcutaneous immunization on days 0, 14 and 28, intravenous boosting on day 42, and splenocyte collection and NS-1 fusion on day 45. Initial screening used wild-type ASGPR1 extracellular protein in ELISA, followed by HepG2 immunofluorescence. The central plots and table summarize antibody titers and clone characterization. The lower immunofluorescence images show HepG2 cells for J31–J37.

### 2.2 Chimeric J31 retains receptor binding and supports selection for bispecific engineering

CDR comparison distinguished candidates for recombinant expression while identifying a closely related group comprising J33, J34 and J36 (Fig. 3A). J34 and J36 shared all six CDR sequences, and J33 shared their heavy-chain CDRs. We selected J34 as a representative of this group and advanced J31, J32, J34, J35 and J37 for chimeric antibody expression.

**Figure 3.**
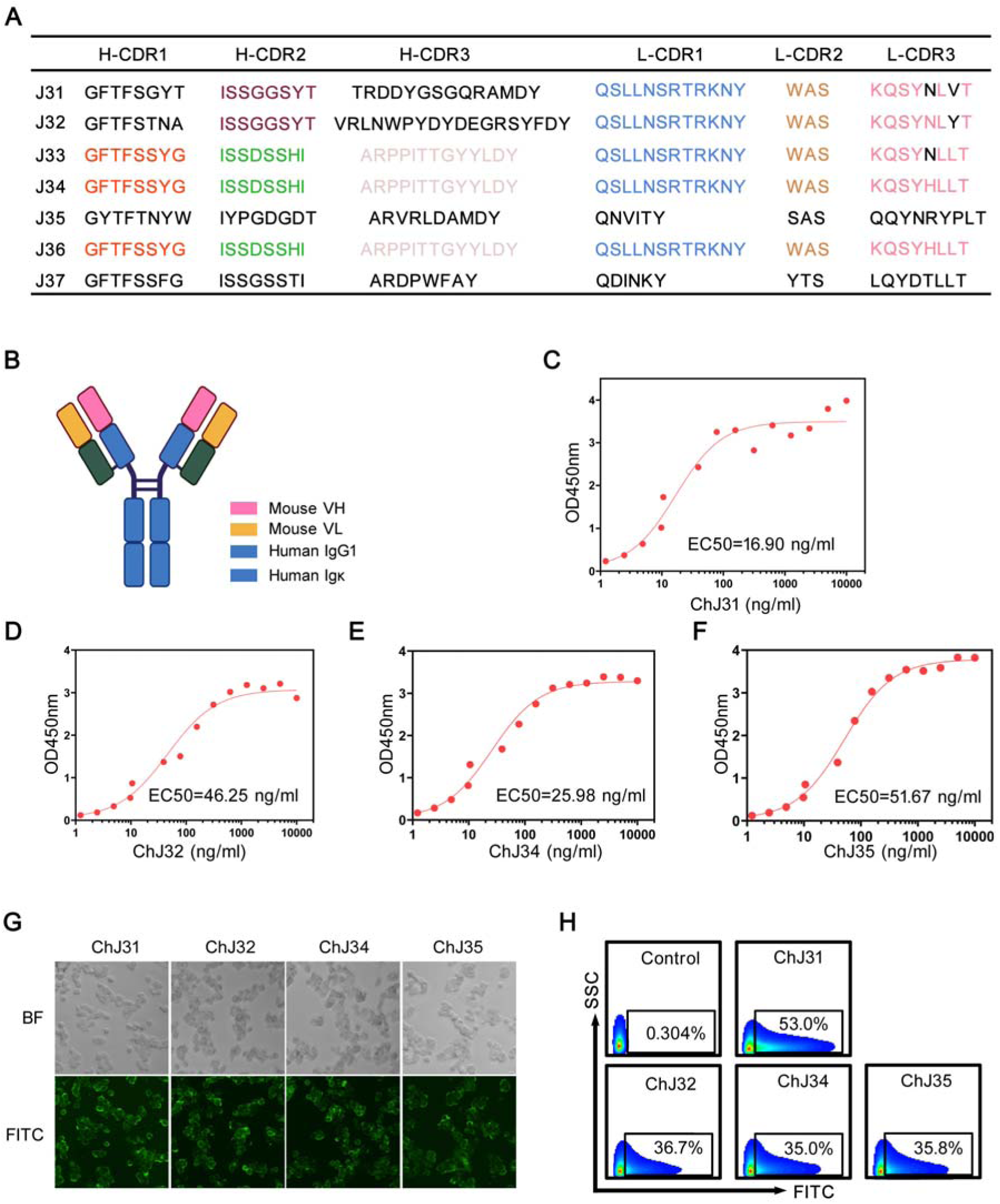
Sequence comparison and characterization of chimeric antibodies. (A) CDR sequence comparison of J31–J37. (B) Mouse–human chimeric antibody design. (C–F) Binding ELISAs for ChJ31, ChJ32, ChJ34 and ChJ35, respectively, using wild-type ASGPR1 (1 μg/mL)-coated wells and HRP-conjugated anti-human IgG (1:5000) detection. EC50 values describe half-maximal assay responses. (G) HepG2 immunofluorescence. (H) Representative flow-cytometry binding plots; displayed percentages indicate positive cells. Quantitative experiments: n = 3 independent experiments; error bars represent SD. Flow analysis used debris exclusion followed by singlet gating, with positivity defined using a human IgG1 isotype control.

Mouse VH and VL domains were joined to human IgG1 heavy-chain and human κ light-chain constant regions (Fig. 3B). ChJ31, ChJ32, ChJ34 and ChJ35 were recovered and characterized by SDS-PAGE (Fig. S1); ChJ37 was not detected in the expression supernatant. All four recovered antibodies showed concentration-dependent binding to immobilized wild-type ASGPR1 (Fig. 3C–F). Their ELISA half-maximal response concentrations were 16.90, 46.25, 25.98 and 51.67 ng/mL, respectively. ChJ31 therefore had the lowest half-maximal response concentration in this assay panel.

Immunofluorescence and flow cytometry confirmed HepG2 binding by the four chimeric antibodies (Fig. 3G,H). The representative flow plots showed 53.0%, 36.7%, 35.0% and 35.8% antibody-positive cells for ChJ31, ChJ32, ChJ34 and ChJ35, respectively. Addition of GalNAc at 10 or 20 μg/mL produced similar antibody-binding signals to the no-GalNAc condition in the displayed assay (Fig. S2), consistent with J31 binding outside the carbohydrate-recognition domain. Together, these results established a panel of recombinant ASGPR1-binding antibodies for evaluation in the bispecific format.

### 2.3 BiAb-J31 combines cell binding with dual-antigen engagement

We paired each ASGPR1-binding candidate with the complete, unmodified adalimumab VH and VL sequences. The adalimumab variable region sequences were obtained from published patent literature. The IgG-like constructs used knobs-into-holes for heavy-chain heterodimerization and CH1–CL crossover in the ASGPR1 arm for light-chain pairing ^[23,24]^. L234A/L235A/P329G substitutions were incorporated into the Fc on the basis of established effector-attenuation strategies ^[26,27]^ (Fig. 4A).

**Figure 4.**
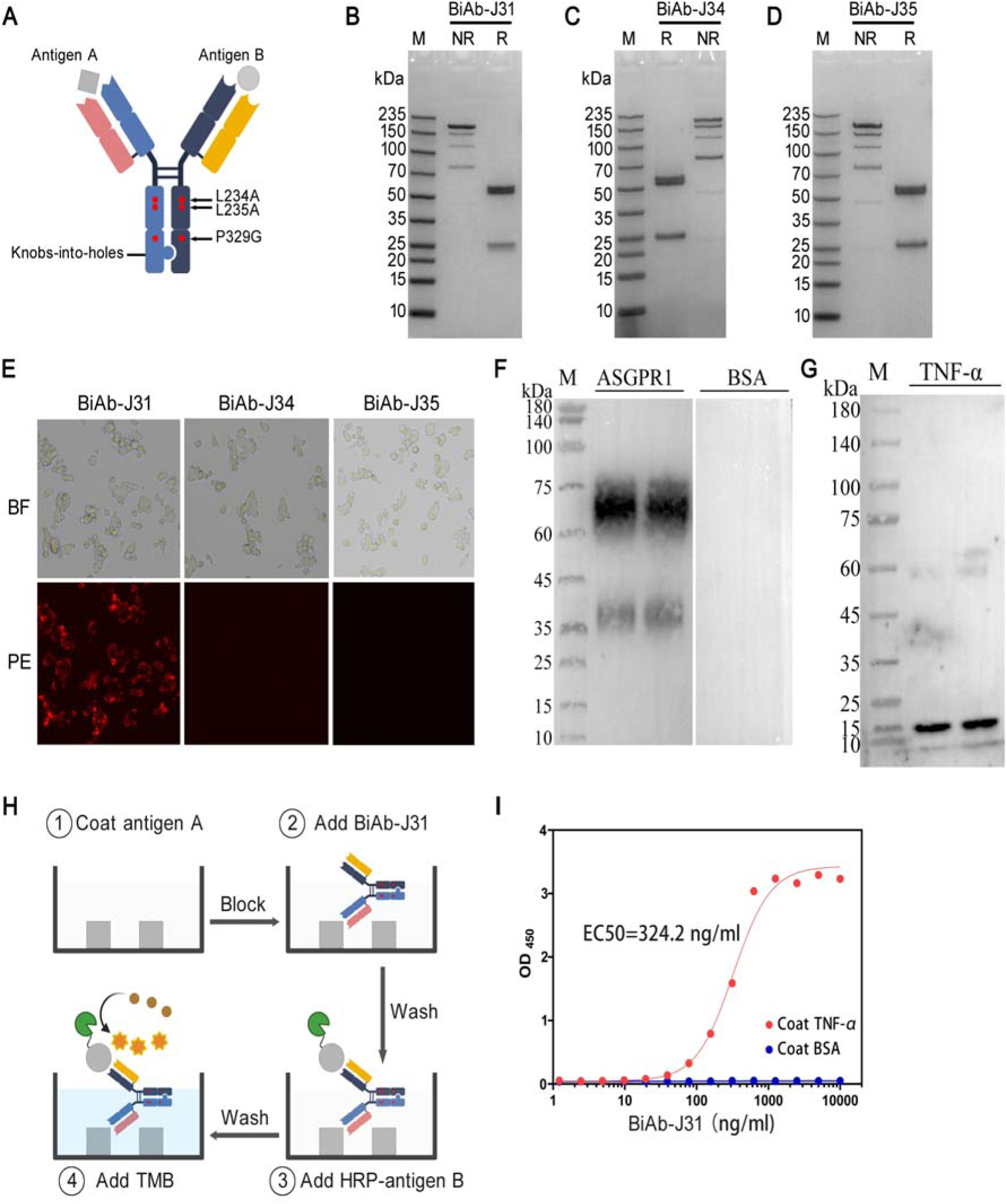
Construction and binding characterization of ASGPR1×TNF-α bispecific antibodies. (A) IgG-like bispecific design with knobs-into-holes engineering, CH1–CL crossover in the ASGPR1 arm, and L234A/L235A/P329G Fc substitutions. (B–D) SDS-PAGE of BiAb-J31, BiAb-J34 and BiAb-J35, respectively. (E) HepG2 binding at 5 μg/mL. (F,G) ASGPR1 and TNF-α immunoblots probed with BiAb-J31 (2 μg/mL) followed by HRP-conjugated anti-human IgG (1:1000). (H) Bridging ELISA design. (I) TNF-α-coated or BSA-coated wells incubated with a BiAb-J31 concentration series followed by HRP-ASGPR1 at 1 μg/mL. EC50 denotes the half-maximal response in this bridging assay. Quantitative experiments: n = 3 independent experiments; error bars represent SD.

BiAb-J31, BiAb-J34 and BiAb-J35 were recovered, whereas BiAb-J32 was not detected in the expression supernatant (Fig. 4B–D). At 5 μg/mL, BiAb-J31 showed detectable HepG2 binding, while BiAb-J34 and BiAb-J35 gave no detectable signal at the concentration examined (Fig. 4E). We therefore selected BiAb-J31 for functional characterization.

BiAb-J31 recognized ASGPR1 and TNF-α in immunoblots (Fig. 4F,G). In a bridging ELISA, TNF-α-bound BiAb-J31 recruited HRP-ASGPR1 in a concentration-dependent manner, with a half-maximal response concentration of 324.2 ng/mL; BSA-coated wells gave background-level signals (Fig. 4H,I). Supplementary assays showed HRP-ASGPR1 binding to immobilized BiAb-J31 and ASGPR1 recruitment after BiAb-J31 binding to membrane-associated TNF-α (Figs. S3,S4). These complementary configurations demonstrate dual-antigen engagement. BiAb-J31 also gave a lower CD64-binding signal than ChJ31 in the supplementary receptor-binding assay (Fig. S5).

### 2.4 BiAb-J31 recruits TNF-α to cells and supports uptake of labeled complexes

We first tested whether cell-associated BiAb-J31 could recruit TNF-α. HepG2 cells were sequentially incubated with BiAb-J31, TNF-α and PE-conjugated anti-TNF-α, with washing between steps (Fig. 5A). The complete sequence produced a cell-associated PE signal, whereas omission conditions and the HEK293T comparison showed no detectable signal in the displayed images (Fig. 5B). This sequential assay demonstrates TNF-α recruitment after BiAb-J31 binding to cells.

**Figure 5.**
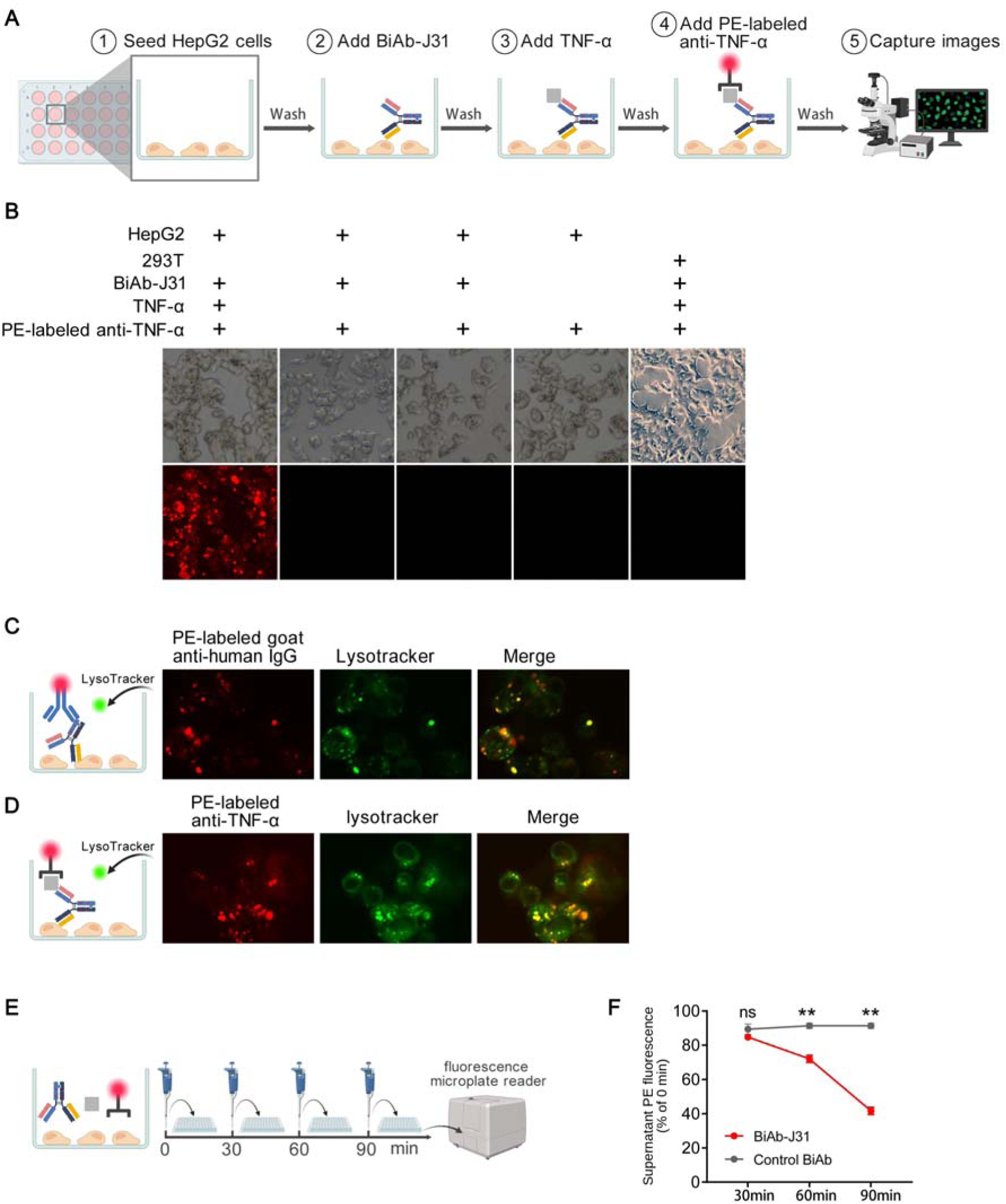
BiAb-J31 supports cellular recruitment and uptake of TNF-α-containing complexes. (A) Sequential recruitment workflow: BiAb-J31, wash, TNF-α, wash, PE-conjugated anti-TNF-α, wash and imaging. (B) Cell-associated PE signal with the indicated omission conditions and HEK293T comparison. (C) Live-cell trafficking of BiAb-J31-containing complexes tracked using PE-conjugated anti-human IgG. (D) Trafficking of TNF-α-containing complexes tracked using PE-conjugated anti-TNF-α. PE detection antibodies were present during the 30-minute uptake period at 37°C in 5% CO₂. LysoTracker (100 nM) was added simultaneously with the bispecific antibody; green fluorescence marks LysoTracker-positive compartments. (E) Supernatant sampling workflow. (F) Relative supernatant PE fluorescence at 0, 30, 60 and 90 minutes after simultaneous addition of bispecific antibody, TNF-α and PE-conjugated anti-TNF-α. The GPC3×CD3 control retains its published Fc configuration ^[28]^. Values are expressed as relative supernatant PE fluorescence (% of time zero). Background was determined using cell culture supernatant without the PE-conjugated detection antibody. Each independent experiment was background-corrected and normalized to the corresponding treatment group at time zero before averaging. Quantitative experiments: n = 3 independent experiments; error bars represent SD. Panels B and C,D were acquired using fluorescence microscopy and Nikon A1 confocal microscopy, respectively. Group comparisons used two-tailed Student’s t tests (GraphPad Prism 5); p < 0.05 was considered significant.

Live-cell imaging then followed uptake using PE-conjugated detection antibodies present during incubation. BiAb-J31-containing complexes tracked with anti-human IgG and TNF-α-containing complexes tracked with anti-TNF-α both produced intracellular PE signals overlapping with LysoTracker-positive compartments (Fig. 5C,D).

To examine the corresponding change in the extracellular compartment, we added BiAb-J31, TNF-α and PE-conjugated anti-TNF-α simultaneously to HepG2 cultures and measured supernatant fluorescence over 90 minutes (Fig. 5E). Relative PE fluorescence fell to approximately 70% of its initial value at 60 minutes and 40% at 90 minutes, while the GPC3×CD3 control bispecific showed substantially less change (Fig. 5F). The control retained its published Fc configuration ^[28]^. PE fluorescence in this assay reports labeled detection-antibody-containing material rather than intact TNF-α directly (see Discussion). Thus, sequential recruitment, intracellular imaging and extracellular reporter loss together support cellular uptake of TNF-α-containing labeled complexes, with an approximately 60% decrease in supernatant PE signal over 90 minutes.

### 2.5 J31 recognition maps to ASGPR1 residues 62–100 near the membrane

To define the receptor region recognized by J31, we generated overlapping ASGPR1 extracellular segments: N, residues 62–151; M, residues 132–221; and C, residues 202–291 (Fig. 6A). Each construct retained the intracellular and transmembrane regions, residues 1–61, and carried an N-terminal eGFP moiety (Fig. 6B). BiAb-J31 bound the N-region construct expressed in HEK293T cells, localizing recognition to the membrane-proximal portion of the extracellular sequence (Fig. 6C).

**Figure 6.**
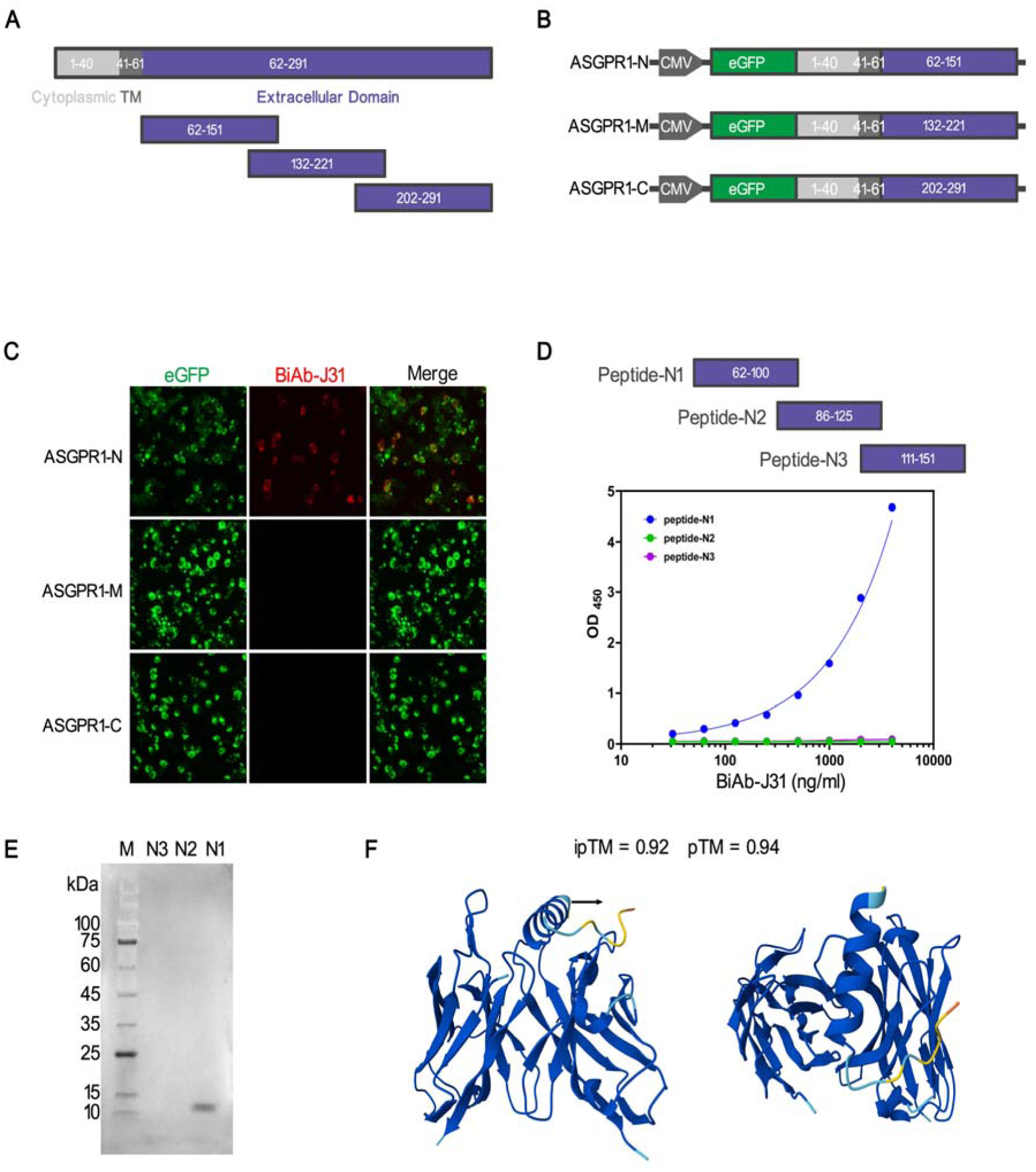
Regional mapping of J31 recognition. (A) ASGPR1 extracellular regions used for mapping. (B) N-terminal eGFP constructs retaining ASGPR1 residues 1–61 and containing N (62–151), M (132–221) or C (202–291). (C) Binding in transfected HEK293T cells. (D) ELISA with free synthetic peptides N1 (62–100), N2 (86–125) and N3 (111–151), each coated at 10 μg/mL, with HRP-conjugated anti-human IgG (1:5000) detection. (E) Synthetic-peptide immunoblot probed with BiAb-J31 (2 μg/mL) followed by HRP-conjugated anti-human IgG (1:1000). (F) AlphaFold 3 model of J31 VH–VL with N1; ipTM = 0.92 and pTM = 0.94. Quantitative experiments: n = 3 independent experiments; error bars represent SD.

Three overlapping synthetic peptides further resolved this region: N1, residues 62–100; N2, residues 86–125; and N3, residues 111–151. N1 showed detectable BiAb-J31 binding in ELISA and immunoblotting, whereas N2 and N3 did not (Fig. 6D,E). The combined cell-fragment and peptide results localize recognition to residues 62–100, identifying a receptor attachment region outside the carbohydrate-recognition domain.

### 2.6 Structural modeling proposes contacts within the J31 Fv–N1 interface

We modeled J31 VH and VL together with the N1 peptide using AlphaFold 3 ^[29]^. The resulting Fv–N1 model had reported confidence scores of ipTM = 0.92 and pTM = 0.94 (Fig. 6F). N1 was positioned at an interface formed by the antibody variable domains, with candidate contacts shown in Fig. 7. The model complements regional mapping by nominating residue-level interactions for subsequent testing.

**Figure 7.**
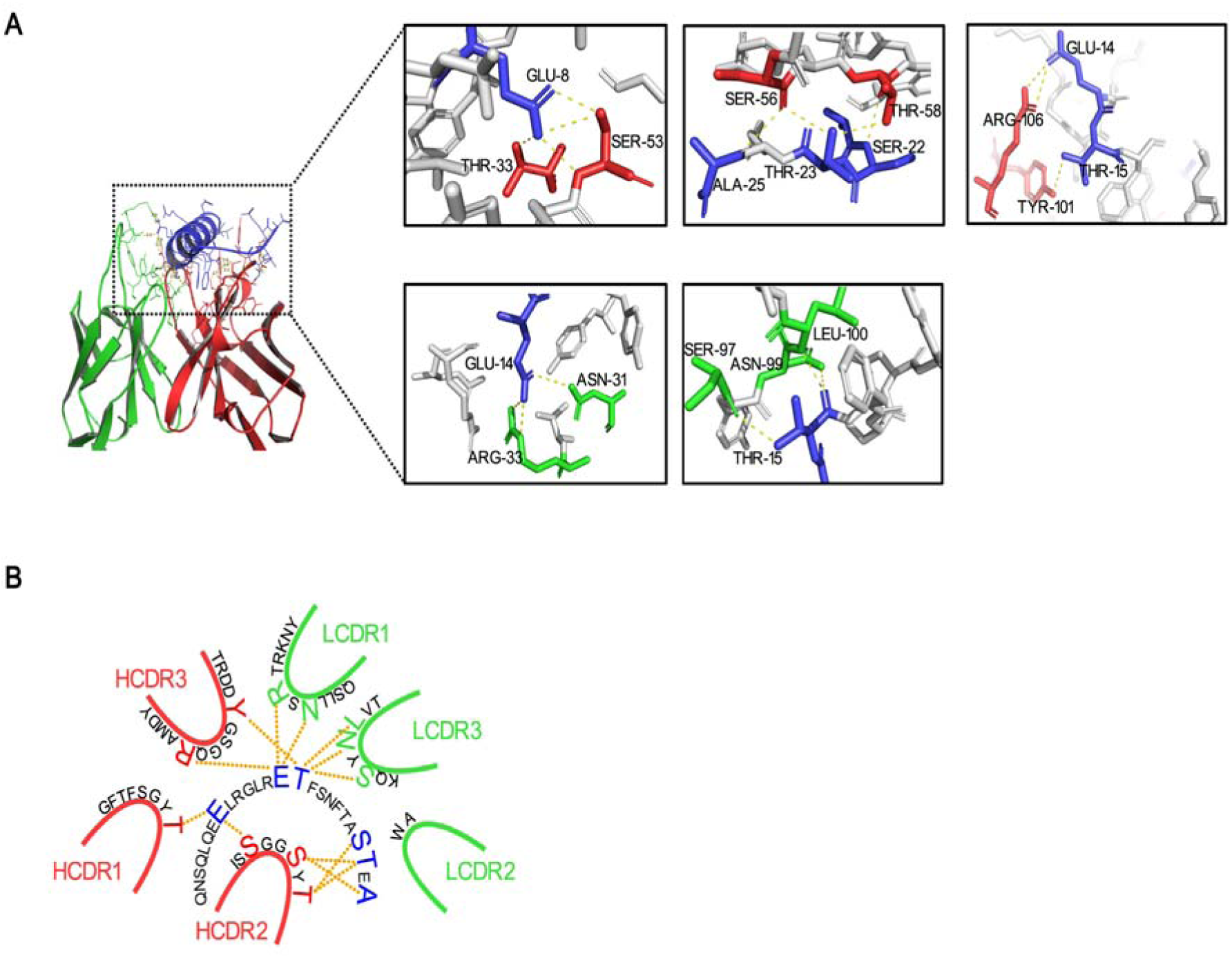
Predicted J31 Fv–N1 interface. (A) Predicted complex and candidate contact views; VH is red, VL is green and N1 is blue. (B) Schematic representation of predicted interactions. The model contains variable domains and N1 only. The displayed contacts are model predictions.

## 3 Discussion

This study identifies a membrane-proximal ASGPR1-binding antibody and connects it to TNF-α recruitment and uptake in an IgG-like bispecific format. The distinguishing molecular feature relative to glycan-based and EndoTag recruiters is J31 recognition of ASGPR1 residues 62–100. This region provides an attachment site outside the carbohydrate-recognition domain, and the J31 arm retains detectable cell binding after combination with adalimumab variable regions. Dual-antigen engagement, sequential TNF-α recruitment and uptake of labeled complexes link this receptor-binding feature to a potential cytokine-uptake application.

The work extends an established field of receptor-directed extracellular protein handling. GalNAc-LYTACs, MoDE-As and PCSK9-directed heterobifunctional molecules have demonstrated ASGPR recruitment across different molecular formats ^[10–13]^. Protein-based EndoTags further established alternative receptor-binding sites as a route to endocytosis ^[20]^. Within this landscape, the contribution of BiAb-J31 is the combination of a regionally mapped antibody recruiter with a defined TNF-α-binding arm. The J31 region and IgG-like format provide concrete features for future comparisons of receptor recruitment, cargo geometry and trafficking. Studies using CI-M6PR-directed nanoplatforms and yeast-derived glycans, or transferrin-receptor-directed vesicle and peptide constructs, likewise illustrate how receptor choice and molecular format shape extracellular targeting strategies ^[30–33]^. Their different targets and endpoints make them design precedents rather than quantitative comparators for the present uptake assay.

The discovery route also has practical relevance. Replacing residues 237–266 reduced representation of that sequence in the immunogen, while screening against wild-type ASGPR1 ensured that subsequent selection was directed toward the unmodified receptor. Identification of an antibody recognizing residues 62–100 is consistent with the design objective. Because a parallel wild-type-immunogen group was not included, the outcome establishes a successful route to J31 rather than the comparative efficiency of the immunogen strategy. The resulting regional map is nevertheless useful for describing the molecule independently of how it was discovered.

Recognition outside the carbohydrate-recognition domain motivates examination of how antibody recruitment interacts with native receptor function. The GalNAc assay showed similar chimeric antibody-binding signals across the tested conditions, providing a starting observation for this question. ASGPR physiology includes glycoprotein handling and platelet-linked thrombopoietin regulation ^[15–18]^, so preservation of native substrate uptake remains a functional property to establish. Work on CI-M6PR-directed LYTACs has also highlighted receptor occupancy, recycling and lysosomal maturation as determinants of activity ^[34]^. These findings suggest that attachment-site selection and the trafficking capacity of the recruited receptor should be evaluated together.

Format conversion was a meaningful selection step. Although four chimeric antibodies bound ASGPR1 and HepG2 cells, only BiAb-J31 retained detectable HepG2 binding among the three recovered bispecific constructs at the concentration examined. This result illustrates the need to reassess receptor-binding antibodies after changes in valency and domain arrangement. Knobs-into-holes and CrossMab address chain pairing ^[22–25]^, but compatibility with the final geometry must still be assessed for each candidate. The reduced CD64-binding signal is consistent with the intended Fc attenuation ^[26,27]^; functional effector activity and pharmacokinetics are distinct properties of the final molecule. In particular, FcRn-dependent IgG recycling provides relevant context for exposure ^[35]^, without determining the persistence of a receptor-targeting bispecific on its own.

The functional assays resolve successive steps in the proposed route. The bridging assays establish dual-antigen engagement, and sequential incubation shows that cell-associated BiAb-J31 can subsequently recruit TNF-α. Imaging and supernatant measurements then support redistribution of fluorescent complexes from the extracellular medium into cells. Interpretation of this last step is bounded by the reporter system: detection antibodies were present during uptake and could influence complex valency, while PE fluorescence reports labeled material rather than an independent measurement of intact TNF-α. LysoTracker overlap supports delivery to acidic compartments but does not establish proteolysis. Moreover, the GPC3×CD3 control differs in target specificity and Fc configuration, so it does not isolate ASGPR1 dependence. The current evidence therefore supports TNF-α recruitment and uptake in the tested complex system; direct cargo measurements and receptor-specific controls would establish how this behavior translates to intact-cytokine depletion.

Regional mapping adds a molecular basis for further development. The overlapping fragments and N1 peptide locate recognition to residues 62–100, while the Fv–N1 prediction supplies candidate contacts within that region. Advances in protein modeling make such hypotheses increasingly accessible ^[29,36]^, although local interface accuracy remains an experimental question. Published assessments in ligand-bound GPCR systems illustrate the broader distinction between overall model quality and local interaction accuracy ^[37]^. Here, the experimentally mapped region carries the principal structural conclusion; the atomic contacts remain predictions. Humanization methods offer a further route for engineering the J31 arm, which currently retains mouse variable domains ^[38]^.

For TNF-α targeting, the use of unmodified adalimumab variable regions provides a defined binding component with an established therapeutic lineage ^[2,3]^. Its incorporation into BiAb-J31 connects cytokine recognition to cell recruitment, but the biological activity of the final construct must be evaluated in its own right. The clinically observed variation in anti-TNF responses provides motivation for exploring additional modes of cytokine control ^[4–6]^, rather than evidence that uptake will overcome treatment resistance. The immediate value of this study is a characterized molecular candidate: J31 supplies a membrane-proximal ASGPR1 attachment site, and the bispecific supports recruitment and uptake of TNF-α-containing complexes. Priority experiments for further development include ASGPR1 loss-of-function controls to confirm receptor dependence, direct cytokine measurements to verify intact TNF-α depletion, and in vivo characterization to assess pharmacokinetics and pharmacodynamic activity. This combination provides a foundation for developing antibody-based extracellular cytokine-uptake strategies.

## 4 Materials and methods

### 4.1 Reagents and cell culture

HepG2 cells were used for antibody-binding, recruitment and trafficking experiments. Huh7 cells were used in hybridoma binding characterization, HEK293T cells served in the indicated comparison and transfection experiments, and NS-1 cells were used for hybridoma generation. The HepG2 uptake assay used DMEM supplemented with 10% fetal bovine serum at 37°C in 5% CO₂, as detailed below.

### 4.2 Immunogen design and preparation

The ASGPR1 extracellular-domain immunogen contained residues 62–291, with Arg237–Asp266 replaced by (G4S)3 and a C-terminal 10×His tag. Structural context was examined using PDB entry 6YAU ^[39]^. This entry contains the GalNAc-derived ligand GN-A. Expression supernatant and nickel-affinity-purified material were analyzed by SDS-PAGE. The immunogen was expressed in HEK293T cells using the pcDNA3.4 vector and PEI-mediated transfection at a PEI-to-total-DNA mass ratio of 3:1. Culture supernatants were collected 72 hours after transfection, and the His-tagged immunogen was purified by nickel-affinity chromatography.

### 4.3 Immunization and hybridoma selection

Female BALB/c mice aged 6–8 weeks were used for immunization. Each mouse received 100 μg of antigen with complete Freund adjuvant subcutaneously on day 0, followed by 50 μg of antigen with incomplete Freund adjuvant subcutaneously on days 14 and 28. An intravenous boost of 50 μg of antigen per mouse was administered on day 42. Splenocytes collected on day 45 were fused with NS-1 myeloma cells using 50% polyethylene glycol 4000 (PEG 4000). Following fusion, cells were selected in hypoxanthine–aminopterin–thymidine (HAT) medium. Hybridomas were first screened using ELISA plates coated with wild-type ASGPR1 extracellular protein and subsequently by HepG2 immunofluorescence. Positive hybridomas underwent three rounds of subcloning by limiting dilution. The resulting clones were designated J31–J37.

### 4.4 Chimeric and bispecific antibody production

Complete mouse VH and VL sequences were fused to human IgG1 heavy-chain and human κ light-chain constant regions. J31, J32, J34, J35 and J37 were subjected to chimeric expression attempts. Both chimeric and bispecific antibodies were expressed in HEK293T cells using pcDNA3.4 vectors and PEI-mediated transfection. PEI and total plasmid DNA were used at a 3:1 mass ratio. For chimeric antibody expression, heavy- and light-chain plasmids were mixed at a 2:1 molar ratio. For bispecific antibody expression, plasmids encoding the ASGPR1 heavy chain, TNF-α heavy chain, ASGPR1 light chain and TNF-α light chain were mixed at a 2:2:1:1 molar ratio. Culture supernatants were collected 72 hours after transfection, and antibodies were purified by Protein A affinity chromatography. Antibodies were eluted with 0.1 M citric acid–sodium citrate buffer (pH 3.2), immediately neutralized with Tris-HCl, and buffer-exchanged into PBS.

For bispecific construction, ASGPR1-binding variable regions were combined with the complete, unmodified adalimumab VH and VL sequences, which were obtained from published patent literature. Knobs-into-holes engineering ^[23]^ introduced T366W into the ASGPR1 heavy chain and T366S/L368A/Y407V into the TNF-α heavy chain. CH1–CL exchange was implemented in the ASGPR1 arm ^[24]^, and L234A/L235A/P329G Fc substitutions were included ^[26]^. BiAb-J31, BiAb-J32, BiAb-J34 and BiAb-J35 were subjected to expression attempts.

The GPC3×CD3 control bispecific antibody was the previously published construct ^[28]^, retaining its original Fc configuration. It was not modified to match the L234A/L235A/P329G Fc of BiAb-J31.

### 4.5 Antigen-binding and bridging ELISAs

For serum-titer determination, hybridoma screening and chimeric antibody-binding ELISAs, wild-type ASGPR1 extracellular protein was coated at 1 μg/mL. For ELISA, coating solutions were prepared in PBS and added at 100 μL per well, followed by overnight incubation at 4°C. Wells were blocked with 5% skim milk powder in PBS for 1 hour at 37°C. The test-antibody incubation was performed for 1 hour at 37°C. Incubation with HRP-conjugated secondary antibodies or HRP-labeled antigens was performed for 30 minutes at 37°C. Between assay incubation steps, wells were washed five times with PBS containing 0.1% Tween-20 (PBST), for 1 minute per wash. Color was developed by adding 100 μL of TMB substrate per well and incubating for 15 minutes at room temperature in the dark. The reaction was stopped by adding 50 μL of 2 M H₂SO₄ per well, and absorbance was measured at 450 nm using a Varioskan LUX multimode microplate reader (Thermo Scientific; Type 3020).

For serum-titer determination and hybridoma screening, bound mouse antibodies were detected using an HRP-conjugated anti-mouse secondary antibody at a dilution of 1:20000.

For chimeric antibody-binding assays, plates were coated with wild-type ASGPR1 extracellular protein, incubated with antibody concentration series, and developed using HRP-conjugated anti-human IgG (1:5000) and TMB. For the GalNAc experiment, ASGPR1-coated wells received fixed antibody concentrations together with 0, 10 or 20 μg/mL GalNAc. ChJ31, ChJ34 and ChJ35 were each used at 0.1 μg/mL and ChJ32 at 0.2 μg/mL. Bound antibody was detected using HRP-conjugated anti-human IgG (1:5000).

For the TNF-α bridging ELISA, wells were coated with TNF-α or BSA at 1 μg/mL, incubated with a BiAb-J31 concentration series and then with HRP-ASGPR1 at 1 μg/mL before TMB development. In the supplementary ASGPR1-binding assay, BiAb-J31 or the GPC3×CD3 control was immobilized at 1 μg/mL and exposed to an HRP-ASGPR1 concentration series. For CD64 binding, wells coated with CD64 at 1 μg/mL received a ChJ31 or BiAb-J31 concentration series followed by HRP-ASGPR1 at 1 μg/mL.

EC50 values for the chimeric antibody-binding assays (Fig. 3C–F) and the TNF-α bridging ELISA (Fig. 4I) were calculated by four-parameter logistic (4PL) regression using GraphPad Prism 9.0 and represent half-maximal responses in the respective immobilized-antigen assays.

### 4.6 Immunoblotting

ASGPR1 and TNF-α immunoblots in Fig. 4F,G were probed with BiAb-J31 (2 μg/mL) followed by HRP-conjugated anti-human IgG (1:1000); BSA served as the indicated negative control. For the supplementary bridging immunoblot, membrane-associated TNF-α was incubated with BiAb-J31 (2 μg/mL) and subsequently HRP-ASGPR1 (2 μg/mL). This bridging detection did not use an HRP-conjugated secondary antibody. Synthetic peptide immunoblots (Fig. 6E) were probed with BiAb-J31 (2 μg/mL) followed by HRP-conjugated anti-human IgG (1:1000).

For these immunoblots (Figs. 4F,G, 6E and S4), samples were transferred by wet transfer onto polyvinylidene difluoride (PVDF) membranes with a pore size of 0.45 μm. Membranes were blocked overnight at 4°C with 5% skim milk powder prepared in PBST. The same blocking solution was used for washing, with four washes of 5 minutes each. BiAb-J31 was incubated for 1 hour at room temperature. Subsequent incubation with HRP-conjugated anti-human IgG (Figs. 4F,G and 6E) or HRP-ASGPR1 (Fig. S4) was performed for 45 minutes at room temperature.

### 4.7 Cell binding and sequential TNF-α recruitment

Hybridoma and chimeric antibody binding were examined by fluorescence microscopy, with chimeric antibody binding additionally assessed using a BD SYMPHONY flow cytometer (BD Biosciences). For flow-cytometric staining, HepG2 cells were incubated separately with ChJ31, ChJ32, ChJ34 or ChJ35 at 2 μg/mL for 20 minutes at 4°C. FITC-conjugated anti-human IgG (1:50) was used for secondary staining for 20 minutes at 4°C. After both primary and secondary staining, cells were washed three times with PBS containing 1% BSA, for 1 minute per wash. A total of 100,000 cells were acquired per sample, and data were analyzed using FlowJo. Debris was excluded using forward- and side-scatter characteristics, followed by singlet gating before determining the percentage of antibody-positive cells. No viability dye was used. The threshold for antibody-binding positivity was set using a commercial human IgG1 isotype control at 2 μg/mL. BiAb-J31, BiAb-J34 and BiAb-J35 were compared on HepG2 cells at 5 μg/mL by fluorescence microscopy.

For sequential recruitment, cells were neither fixed nor permeabilized, to avoid potential fixation-induced changes to membrane antigens. Cells were incubated with BiAb-J31 (500 ng/mL) for 30 minutes at 4°C, washed, incubated with TNF-α (100 ng/mL) for 30 minutes at 4°C, washed, and incubated with PE-conjugated anti-TNF-α (1:50) for 30 minutes at 4°C before a final wash and imaging. After each incubation, cells were washed three times with sterile PBS for 1 minute per wash. Images were acquired in sterile PBS containing 1% BSA using a fluorescence microscope. Omission controls and HEK293T comparisons were included as displayed.

### 4.8 Live-cell trafficking and supernatant fluorescence assay

BiAb-J31 was used at a final concentration of 100 ng/mL in both imaging conditions. PE-conjugated anti-human IgG (final dilution, 1:100) was used to track BiAb-J31-containing complexes in Fig. 5C. In Fig. 5D, TNF-α was used at a final concentration of 30 ng/mL, and PE-conjugated anti-TNF-α (final dilution, 1:50) was used to track TNF-α-containing complexes. For Fig. 5D, BiAb-J31, TNF-α and PE-conjugated anti-TNF-α were added directly and simultaneously to the cells, without preincubation outside the cell culture. The detection antibodies were present during live-cell uptake. LysoTracker was added simultaneously with the bispecific antibody at a final concentration of 100 nM. Cells were incubated for 30 minutes at 37°C in 5% CO₂, washed three times with sterile PBS for 1 minute per wash, and imaged in sterile PBS containing 1% BSA using a Nikon A1 confocal microscope (Nikon).

HepG2 cells were seeded in 24-well plates at 1 × 10⁵ cells per well and cultured overnight. BiAb-J31 or the GPC3×CD3 control bispecific antibody (final concentration, 100 ng/mL), TNF-α (final concentration, 10 ng/mL), and PE-conjugated anti-TNF-α antibody (final dilution, 1:100) were added simultaneously. Supernatant samples (100 μL) were collected at 0, 30, 60 and 90 minutes and transferred to a black 96-well plate. PE fluorescence was measured using a Varioskan LUX multimode microplate reader (Thermo Scientific; Type 3020) with excitation at 496 nm and emission at 578 nm. Cell culture supernatant prepared without the PE-conjugated detection antibody was used to determine background fluorescence. Background subtraction and normalization to the corresponding treatment group’s time-zero value were performed separately for each of three independent experiments. The normalized values were then averaged and expressed as relative supernatant PE fluorescence (% of time zero).

### 4.9 Regional epitope mapping

The N, M and C extracellular fragments comprised ASGPR1 residues 62–151, 132–221 and 202–291, respectively. Each construct retained residues 1–61 and an N-terminal eGFP moiety. Transfected HEK293T cells were examined by fluorescence microscopy. Overlapping peptides N1 (62–100), N2 (86–125) and N3 (111–151) were chemically synthesized as free peptides and evaluated by ELISA and BiAb-J31 immunoblotting. For the peptide ELISA (Fig. 6D), N1, N2 and N3 were each coated at 10 μg/mL. Bound BiAb-J31 was detected using HRP-conjugated anti-human IgG at a dilution of 1:5000.

### 4.10 Structural modeling

J31 VH, J31 VL and the N1 peptide were supplied to AlphaFold 3 ^[29]^. No CH1 or CL domains were included. The displayed model had ipTM and pTM values of 0.92 and 0.94, respectively.

### 4.11 Replication and statistics

Quantitative experiments were performed in three independent experiments. Error bars represent standard deviation (SD). All statistical tests were performed in GraphPad Prism 5 (GraphPad Software, La Jolla, CA, USA). Comparisons between groups were performed using two-tailed Student’s t tests, with p < 0.05 considered statistically significant. EC50 values for Fig. 3C–F and Fig. 4I were estimated separately using four-parameter logistic regression in GraphPad Prism 9.0 (GraphPad Software, San Diego, CA, USA). Flow-cytometry percentages are values from representative plots.

## Declarations

Animal experiments were approved by the Laboratory Animal Ethics Committee of Southern Medical University.

### Funding

This work was supported by the National Natural Science Foundation of China (Grant No. 32370986).

### Competing interests

Jiang XiaoTao, Li SiJie, Song Xu and Liu Xuan are named inventors on patent application No. 2025114904635, filed by Southern Medical University, relating to the antibodies described in this study. The authors declare no other competing interests.

### Data availability

The underlying data supporting the findings of this study are available from the corresponding authors upon reasonable request.

### Author contributions

Song Xu performed the majority of the experiments. Jiang XiaoTao and Liu Xuan were primarily responsible for experimental design. Jiang XiaoTao and Aojiasi Hashan were primarily responsible for writing the manuscript. Li SiJie, Du JianMing and Sun YiTong contributed to the experimental work.

## Supplementary materials

***Figure S1. Chimeric antibody characterization.** (A–D) SDS-PAGE of ChJ31, ChJ32, ChJ34 and ChJ35, respectively. M, molecular-mass marker; NR, non-reducing; R, reducing. Lane order is M/NR/R in A and D and M/R/NR in B and C. Molecular masses are indicated in kDa.*

***Figure S2. Effect of GalNAc on chimeric antibody binding to ASGPR1.** Wild-type ASGPR1-coated wells received fixed chimeric antibody concentrations together with GalNAc at 0, 10 or 20 μg/mL. ChJ31, ChJ34 and ChJ35 were used at 0.1 μg/mL and ChJ32 at 0.2 μg/mL. Bound antibody was detected using HRP-conjugated anti-human IgG (1:5000). Quantitative experiments: n = 3 independent experiments; error bars represent SD. Group comparisons used two-tailed Student’s t tests (GraphPad Prism 5); p < 0.05 was considered significant.*

***Figure S3. ASGPR1 binding to immobilized bispecific antibodies.** BiAb-J31 or the published GPC3×CD3 control bispecific antibody ^[28]^ was immobilized at 1 μg/mL and incubated with an HRP-ASGPR1 concentration series before TMB development. The concentration gradient is HRP-ASGPR1, not bispecific antibody. Quantitative experiments: n = 3 independent experiments; error bars represent SD.*

***Figure S4. Bridging immunoblot with TNF-α and ASGPR1.** Membrane-associated TNF-α was incubated with BiAb-J31 (2 μg/mL) and then HRP-ASGPR1 (2 μg/mL). Detection therefore reports ASGPR1 recruitment under this membrane-based binding configuration and does not use an HRP-conjugated secondary antibody.*

***Figure S5. CD64-binding assay.** Wells coated with CD64 at 1 μg/mL were incubated with ChJ31 or BiAb-J31 concentration series and then HRP-ASGPR1 at 1 μg/mL before development. The assay compares CD64-binding signals in the indicated detection format. Quantitative experiments: n = 3 independent experiments; error bars represent SD.*

## Notes

### Competing Interest Statement

The authors have declared no competing interest.

## References

1. Feldmann M, Maini RN. Anti-TNF Therapy, from Rationale to Standard of Care: What Lessons Has It Taught Us. The Journal of Immunology. 2010;185:791–794. 10.4049/jimmunol.1090051

2. Tracey D, Klareskog L, Sasso EH, Salfeld JG, Tak PP. Tumor necrosis factor antagonist mechanisms of action: A comprehensive review. Pharmacology & Therapeutics. 2008;117:244–279. 10.1016/j.pharmthera.2007.10.001

3. van de Putte LBA, Atkins C, Malaise M, Sany J, Russell AS, van Riel PLCM, et al. Efficacy and safety of adalimumab as monotherapy in patients with rheumatoid arthritis for whom previous disease modifying antirheumatic drug treatment has failed. Annals of the Rheumatic Diseases. 2004;63:508–516. 10.1136/ard.2003.013052

4. Liang Y, Li Y, Lee C, Yu Z, Chen C, Liang C. Ulcerative colitis: molecular insights and intervention therapy. Molecular Biomedicine. 2024;5:42. 10.1186/s43556-024-00207-w

5. Kumar M, Murugesan S, Ibrahim N, Elawad M, Al Khodor S. Predictive biomarkers for anti-TNF alpha therapy in IBD patients. Journal of Translational Medicine. 2024;22:284. 10.1186/s12967-024-05058-1

6. Kapizioni C, Desoki R, Lam D, Balendran K, Al-Sulais E, Subramanian S, et al. Biologic Therapy for Inflammatory Bowel Disease: Real-World Comparative Effectiveness and Impact of Drug Sequencing in 13 222 Patients within the UK IBD BioResource. Journal of Crohn’s and Colitis. 2024;18:790–800. 10.1093/ecco-jcc/jjad203

7. Zhao L, Zhao J, Zhong K, Tong A, Jia D. Targeted protein degradation: mechanisms, strategies and application. Signal Transduction and Targeted Therapy. 2022;7:113. 10.1038/s41392-022-00966-4

8. Zhang C, Liu Y, Li G, Yang Z, Han C, Sun X, et al. Targeting the undruggables—the power of protein degraders. Science Bulletin. 2024;69:1776–1797. 10.1016/j.scib.2024.03.056

9. Banik SM, Pedram K, Wisnovsky S, Ahn G, Riley NM, Bertozzi CR. Lysosome-targeting chimaeras for degradation of extracellular proteins. Nature. 2020;584:291–297. 10.1038/s41586-020-2545-9

10. Ahn G, Banik SM, Miller CL, Riley NM, Cochran JR, Bertozzi CR. LYTACs that engage the asialoglycoprotein receptor for targeted protein degradation. Nature Chemical Biology. 2021;17:937–946. 10.1038/s41589-021-00770-1

11. Zhou Y, Teng P, Montgomery NT, Li X, Tang W. Development of Triantennary N-Acetylgalactosamine Conjugates as Degraders for Extracellular Proteins. ACS Central Science. 2021;7:499–506. 10.1021/acscentsci.1c00146

12. Caianiello DF, Zhang M, Ray JD, Howell RA, Swartzel JC, Branham EMJ, et al. Bifunctional small molecules that mediate the degradation of extracellular proteins. Nature Chemical Biology. 2021;17:947–953. 10.1038/s41589-021-00851-1

13. Bagdanoff JT, Smith TM, Allan M, O’Donnell P, Nguyen Z, Moore EA, et al. Clearance of plasma PCSK9 via the asialoglycoprotein receptor mediated by heterobifunctional ligands. Cell Chemical Biology. 2023;30:97–109.e9. 10.1016/j.chembiol.2022.12.003

14. Li YY, Yang Y, Zhang RS, Ge RX, Xie SB. Targeted degradation of membrane and extracellular proteins with LYTACs. Acta Pharmacologica Sinica. 2025;46:1–7. 10.1038/s41401-024-01364-y

15. Ashwell G, Morell AG. The Role of Surface Carbohydrates in the Hepatic Recognition and Transport of Circulating Glycoproteins. Advances in Enzymology - and Related Areas of Molecular Biology. 1974;41:99–128. 10.1002/9780470122860.ch3

16. Stockert RJ. The asialoglycoprotein receptor: relationships between structure, function, and expression. Physiological Reviews. 1995;75:591–609. 10.1152/physrev.1995.75.3.591

17. Grewal PK. The Ashwell–Morell Receptor. Methods in Enzymology. 2010;479:223–241. 10.1016/s0076-6879(10)79013-3

18. Grozovsky R, Begonja AJ, Liu K, Visner G, Hartwig JH, Falet H, et al. The Ashwell-Morell receptor regulates hepatic thrombopoietin production via JAK2-STAT3 signaling. Nature Medicine. 2015;21:47–54. 10.1038/nm.3770

19. Meier M, Bider MD, Malashkevich VN, Spiess M, Burkhard P. Crystal Structure of the Carbohydrate Recognition Domain of the H1 Subunit of the Asialoglycoprotein Receptor. Journal of Molecular Biology. 2000;300:857–865. 10.1006/jmbi.2000.3853

20. Huang B, Abedi M, Ahn G, Coventry B, Sappington I, Tang C, et al. Designed endocytosis-inducing proteins degrade targets and amplify signals. Nature. 2025;638:796–804. 10.1038/s41586-024-07948-2

21. Labrijn AF, Janmaat ML, Reichert JM, Parren PWHI. Bispecific antibodies: a mechanistic review of the pipeline. Nature Reviews Drug Discovery. 2019;18:585–608. 10.1038/s41573-019-0028-1

22. Merchant AM, Zhu Z, Yuan JQ, Goddard A, Adams CW, Presta LG, et al. An efficient route to human bispecific IgG. Nature Biotechnology. 1998;16:677–681. 10.1038/nbt0798-677

23. Ridgway JB, Presta LG, Carter P. ‘Knobs-into-holes’ engineering of antibody CH3 domains for heavy chain heterodimerization. Protein Engineering. 1996;9:617–621. 10.1093/protein/9.7.617

24. Schaefer W, Regula JT, Bähner M, Schanzer J, Croasdale R, Dürr H, et al. Immunoglobulin domain crossover as a generic approach for the production of bispecific IgG antibodies. Proceedings of the National Academy of Sciences. 2011;108:11187–11192. 10.1073/pnas.1019002108

25. Klein C, Sustmann C, Thomas M, Stubenrauch K, Croasdale R, Schanzer J, et al. Progress in overcoming the chain association issue in bispecific heterodimeric IgG antibodies. mAbs. 2012;4:653–663. 10.4161/mabs.21379

26. Schlothauer T, Herter S, Koller CF, Grau-Richards S, Steinhart V, Spick C, et al. Novel human IgG1 and IgG4 Fc-engineered antibodies with completely abolished immune effector functions. Protein Engineering Design and Selection. 2016;29:457–466. 10.1093/protein/gzw040

27. Lo M, Kim HS, Tong RK, Bainbridge TW, Vernes JM, Zhang Y, et al. Effector-Attenuating Substitutions That Maintain Antibody Stability and Reduce Toxicity in Mice. Journal of Biological Chemistry. 2017;292:3900–3908. 10.1074/jbc.m116.767749

28. Zhou X, Liu Y, Liu X, Song X, Li S, Chen P, et al. Novel GPC3 N-terminal bispecific antibody exhibits dual anti-tumor effect against tumor cells. Investigational New Drugs. 2025;43:588–601. 10.1007/s10637-025-01530-x

29. Abramson J, Adler J, Dunger J, Evans R, Green T, Pritzel A, et al. Accurate structure prediction of biomolecular interactions with AlphaFold 3. Nature. 2024;630:493–500. 10.1038/s41586-024-07487-w

30. Xing Y, Li J, Wang L, Zhu Z, Yan J, Liu Y, et al. A Bifunctional Lysosome-Targeting Chimera Nanoplatform for Tumor-Selective Protein Degradation and Enhanced Cancer Immunotherapy. Advanced Materials. 2025;37:2417942. 10.1002/adma.202417942

31. Kim S, Kang J, An D, Seo J, Oh DB. Lysosome-Targeting Chimera Using Mannose-6-Phosphate Glycans Derived from Glyco-Engineered Yeast. Bioconjugate Chemistry. 2025;36:424–436. 10.1021/acs.bioconjchem.4c00512

32. Su LY, Tian Y, Zheng Q, Cao Y, Yao M, Wang S, et al. Anti-tumor immunotherapy using engineered bacterial outer membrane vesicles fused to lysosome-targeting chimeras mediated by transferrin receptor. Cell Chemical Biology. 2024;31:1219–1230.e5. 10.1016/j.chembiol.2024.01.002

33. Xiao Y, He Z, Li W, Chen D, Niu X, Yang X, et al. A covalent peptide-based lysosome-targeting protein degradation platform for cancer immunotherapy. Nature Communications. 2025;16:1388. 10.1038/s41467-025-56648-6

34. Ahn G, Riley NM, Kamber RA, Wisnovsky S, Moncayo von Hase S, Bassik MC, et al. Elucidating the cellular determinants of targeted membrane protein degradation by lysosome-targeting chimeras. Science. 2023;382:eadf6249. 10.1126/science.adf6249

35. Roopenian DC, Akilesh S. FcRn: the neonatal Fc receptor comes of age. Nature Reviews Immunology. 2007;7:715–725. 10.1038/nri2155

36. Chen L, Li Q, Nasif KFA, Xie Y, Deng B, Niu S, et al. AI-Driven Deep Learning Techniques in Protein Structure Prediction. International Journal of Molecular Sciences. 2024;25:8426. 10.3390/ijms25158426

37. He XH, Li JR, Shen SY, Xu HE. AlphaFold3 versus experimental structures: assessment of the accuracy in ligand-bound G protein-coupled receptors. Acta Pharmacologica Sinica. 2025;46:1111–1122. 10.1038/s41401-024-01429-y

38. Choi Y, Hua C, Sentman CL, Ackerman ME, Bailey-Kellogg C. Antibody humanization by structure-based computational protein design. mAbs. 2015;7:1045–1057. 10.1080/19420862.2015.1076600

39. RCSB Protein Data Bank. Entry 6YAU: Crystal structure of ASGPR1 in complex with GN-A. Released 13 January 2021. 10.2210/pdb6YAU/pdb

