## Supplementary Figures for "A bispecific antibody targeting a membrane-proximal ASGPR1 region promotes TNF-α recruitment and uptake"

#### Figure S1 Chimeric antibody characterization

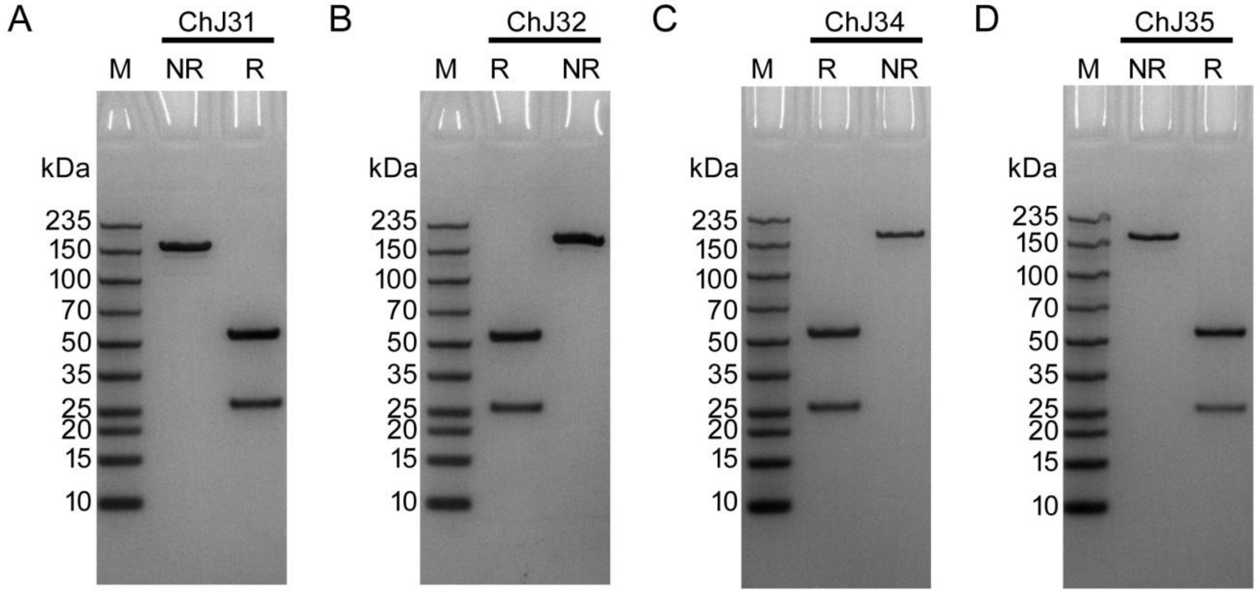


(A–D) SDS-PAGE of ChJ31, ChJ32, ChJ34 and ChJ35, respectively. M, molecular-mass marker; NR, non-reducing; R, reducing. Lane order is M/NR/R in A and D and M/R/NR in B and C. Molecular masses are indicated in kDa.

### Figure S2 Effect of GalNAc on chimeric antibody binding to ASGPR1

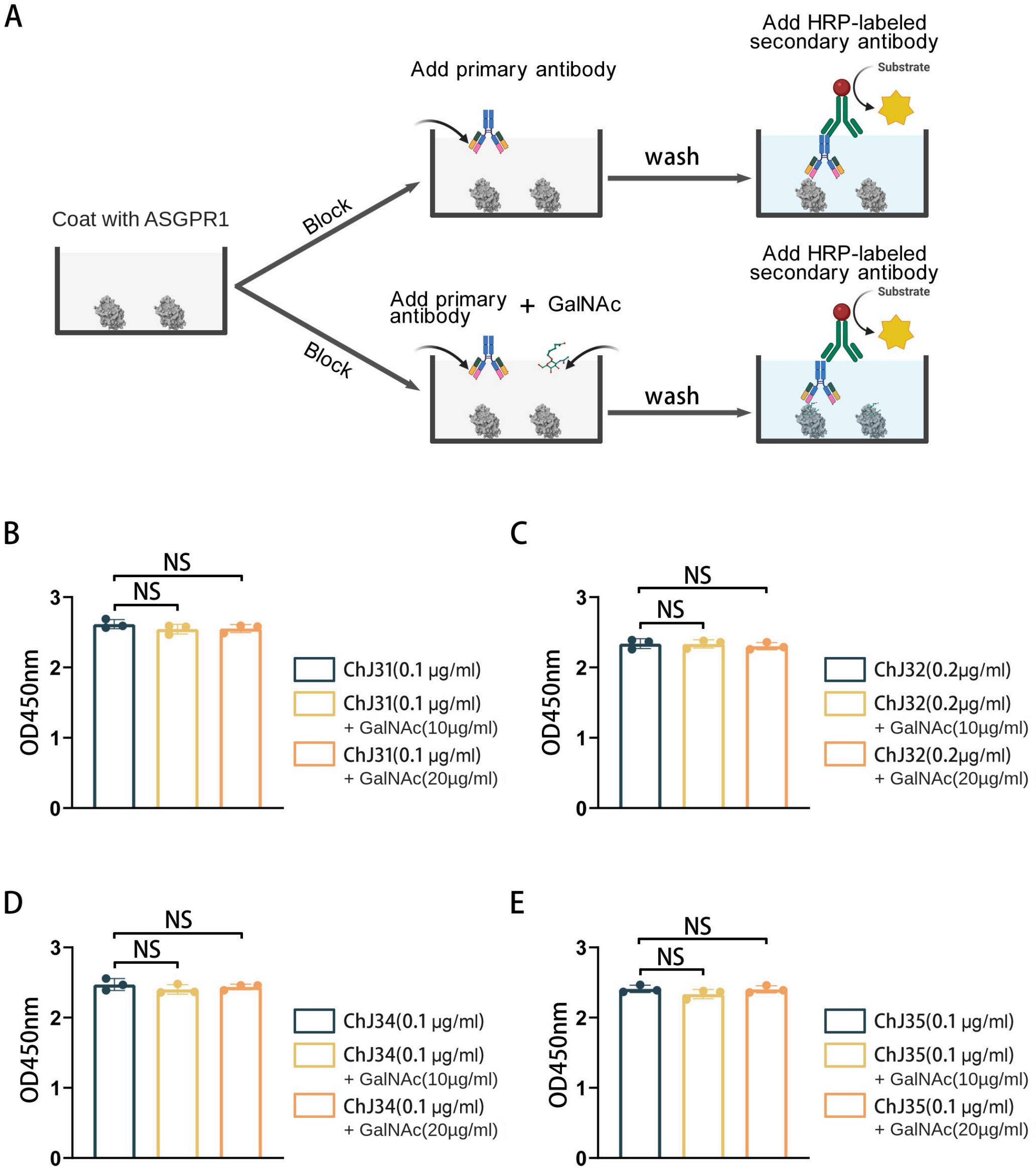


Wild-type ASGPR1-coated wells received fixed chimeric antibody concentrations together with GalNAc at 0, 10 or 20 µg/mL. ChJ31, ChJ34 and ChJ35 were used at 0.1 µg/mL and ChJ32 at 0.2 µg/mL. Bound antibody was detected using HRP-conjugated anti-human IgG (1:5000). Quantitative experiments: n = 3 independent experiments; error bars represent SD. Group comparisons used two-tailed Student's t tests (GraphPad Prism 5); $p < 0.05$ was considered significant.

#### Figure S3 ASGPR1 binding to immobilized bispecific antibodies

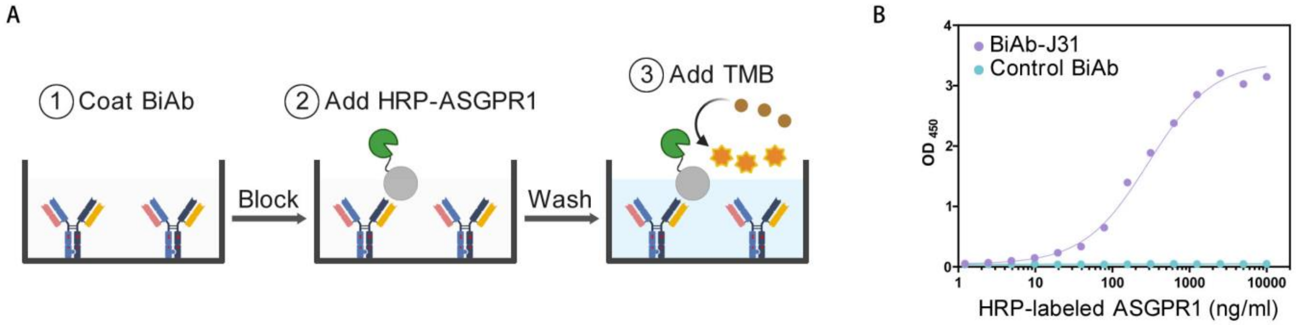


BiAb-J31 or the published GPC3×CD3 control bispecific antibody [28] was immobilized at 1 μg/mL and incubated with an HRP-ASGPR1 concentration series before TMB development. The concentration gradient is HRP-ASGPR1, not bispecific antibody. Quantitative experiments: n = 3 independent experiments; error bars represent SD.

**Figure S4 Bridging immunoblot with TNF-α and ASGPR1**

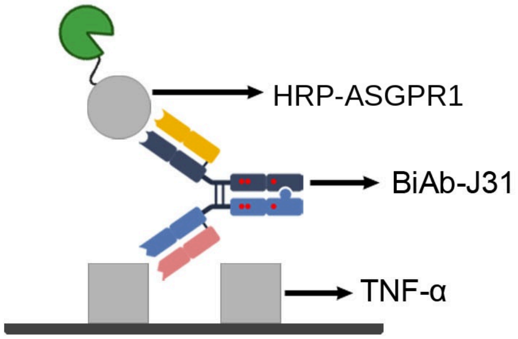


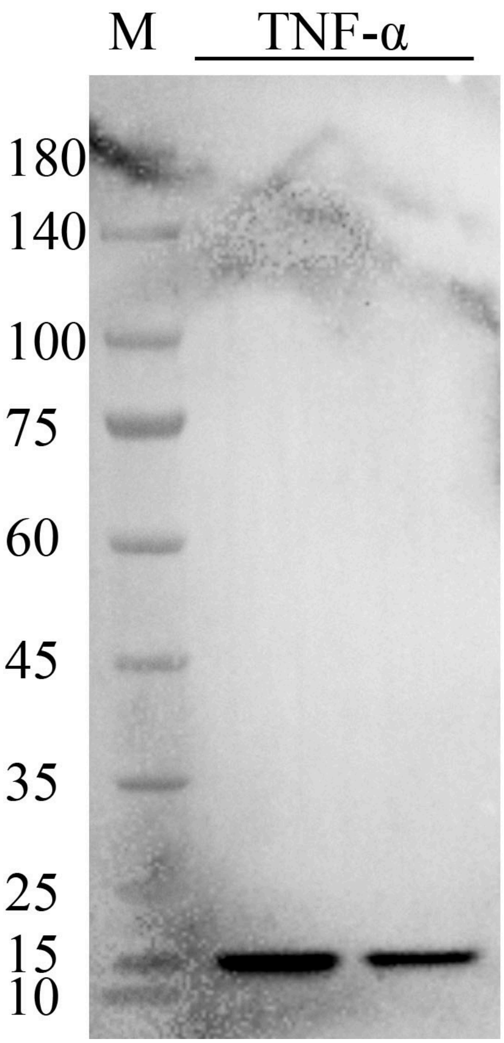


Membrane-associated TNF-α was incubated with BiAb-J31 (2 μg/mL) and then HRP-ASGPR1 (2 μg/mL). Detection therefore reports ASGPR1 recruitment under this membrane-based binding configuration and does not use an HRP-conjugated secondary antibody.

#### Figure S5 CD64-binding assay

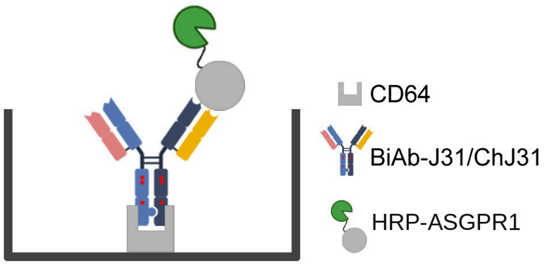


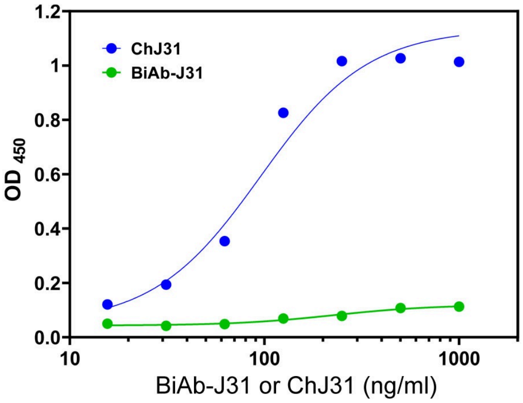


Wells coated with CD64 at 1 μg/mL were incubated with ChJ31 or BiAb-J31 concentration series and then HRP-ASGPR1 at 1 μg/mL before development. The assay compares CD64-binding signals in the indicated detection format. Quantitative experiments: n = 3 independent experiments; error bars represent SD.
